# Leveraging Grating-Based Flickers: A Leap Toward Practical, Visually Comfortable, and High-Performance Dry EEG Code-VEP BCI

**DOI:** 10.1101/2024.07.17.603960

**Authors:** Frédéric Dehais, Kalou Cabrera Castillos, Simon Ladouce, Pierre Clisson

## Abstract

**Purpose:** Reactive Brain-Computer Interfaces (rBCIs) typically rely on repetitive visual stimuli, which can strain the eyes and cause attentional distraction. To address these challenges, we propose a novel approach rooted in visual neuroscience to design visual Stimuli for Augmented Response (StAR). The StAR stimuli consist of small randomly-oriented *Gabor* or *Ricker* patches that optimize foveal neural response while reducing peripheral distraction.

**Methods:** In a factorial design study, 24 participants equipped with an 8-dry electrodes EEG system focused on series of target flickers presented under three formats: traditional ’Plain’ flickers, *Gabor*-based, or *Ricker*-based flickers. These flickers were part of a five-classes Code Visually Evoked Potentials (c-VEP) paradigm featuring low frequency, short, and aperiodic visual flashes.

**Results:** Subjective ratings revealed that *Gabor* and *Ricker* gratings were visually comfortable and nearly invisible in peripheral vision compared to plain flickers. Moreover, *Gabor* and *Ricker*-based textures achieved higher accuracy (93.6% and 96.3%, respectively) with only 88 seconds of calibration data, compared to plain flickers (65.6%). A follow-up online implementation of this experiment was conducted to validate our findings within the frame of naturalistic operations. During this trial, remarkable accuracies of 97.5% in a cued task and 94.3% in an asynchronous digicode task were achieved, with a mean decoding time as low as 1.68 seconds.

**Conclusion:** This work demonstrates the potential to expand BCI applications beyond the lab by integrating visually unobtrusive systems with gel-free, low density EEG technology, thereby making BCIs more accessible and efficient. The datasets, algorithms, and BCI implementations are shared through open-access repositories.

## 1 Introduction

Reactive Brain-Computer Interfaces (rBCIs) based on Visual Evoked Potentials (VEPs) leverage neural responses triggered by the presentation of repeated visual stimuli (RVS), either periodic (such as Steady-States or SSVEP) or aperiodic (code-VEP, C-VEP). These interfaces utilize diverse methods, from basic flickers alternating between contrasting colors to more intricate stimuli such as pattern reversal checker-boards and motion-based animations [1]. Regardless of the chosen approach, RVS have been associated with a range of visual discomfort issues [2–11]. Another critical yet mostly overlooked issue pertains to the simultaneous presentation of multiple flashing stimuli, which can divert the user’s attention away from the intended task that the BCI aims to support or assist with. Furthermore, when users focus on an RVS to input a mental command, the presence of peripheral flickers demands active top-down attentional focus to filter out distractions [12]. Despite the significant cognitive effort deployed to filter out stimuli in the periphery, which can amount to mental fatigue, the EEG signal may still be contaminated by neighboring flickers, potentially hindering classification performance [13, 14].

To address visual discomfort in VEP-based rBCIs, recent research has prioritized on the optimization of user experience while maintaining classification accuracy. Various approaches to the design of RVS stimuli have been explored to improve visual comfort, including the increase of flicker frequency range [15–18], the reduction of contrast between flicker phases [15, 19, 20], and the use of textured stimuli [21, 22]. However, while these approaches hold promise in enhancing the user experience of stimuli presented in central vision, they do not prevent users from being distracted by neighboring stimuli in peripheral vision. To mitigate this effect, the formerly NextMind company pioneered VEP-based stimuli that had then only received limited attention within the rBCI community. Their flickers consisted of small contrasted white bars surrounded by black and greyish borders, forming contours that help attenuate peripheral distraction. However, the system was marketed as a closed system with no access to raw data, lacking any published insights into the design of the stimuli or the underlying classification algorithm. Building on these different previous findings (low amplitude depth, contrasted textures), we are focusing on designing new stimuli called StAR (Stimuli for Augmented Response) to address both foveal visual discomfort and peripheral contamination while maintaining high accuracy.

Our proposition with the StAR stimuli is to incorporate randomly oriented gratings within which we reduce the amplitude depth as previously done. With these gratings, widely employed in visual system studies [23], we generate well-contrasted yet globally low-luminance stimuli that are visually comfortable while eliciting strong VEP responses measurable by surface electrodes [24]. These stimuli effectively tap into the responses of retinal cells, particularly on-off-centered bipolar cells, which play a crucial role in edge and contrast detection [25]. On-off bipolar cells demonstrate heightened sensitivity to changes in luminance, firing both during light increment and decrement [26], thus proving effective at capturing boundaries and transitions between different illumination levels, akin to an edge-enhanced image filter [27]. By introducing diversity in the orientation of the patches, we anticipate bolstering response in the visual pathway by engaging a wider spectrum of orientation-selective neurons, particularly in regions such as the primary visual cortex (V1) [28].

By reducing the size of these gratings and minimizing their amplitude modulation depth, our aim is not only to enhance visual comfort but also to ensure their minimal perceptibility in peripheral vision, thereby reducing the potential for cross-contamination between simultaneously presented flickers. The lower saliency of grating stimuli presented in peripheral vision stems from the difference in sensitivity to low-level visual features between cones and rods photoreceptors [29]. Cone cells, abundant in the macula, exhibit high responsiveness to grating stimuli, resulting in the high visual acuity of the foveal focus. Conversely, rods photoreceptors, primarily distributed in the outer areas of the retina, despite their high sensitivity to luminance changes, demonstrate diminished sensitivity to grating stimuli, leading to the perception of faint, blurry gratings presented in the periphery. Furthermore, the perceptual reduction in peripheral vision results from the presence of numerous small and identical gratings with minimal spacing between them in our StAR stimuli, a phenomenon known as the ”crowding effect [30]. Identifying individual objects in peripheral vision is unproblematic when they are adequately spaced. Conversely, when objects such as gratings are grouped together with high degree of resemblance, the crowding effect impedes the ability to discern them [31].

In this study, our objective is to demonstrate that StAR stimuli not only alleviate foveal and peripheral visual comfort issues but also enhance brain response. We design these stimuli using either *Gabor* filter-based or Ricker wavelet-based patches. While *Gabor* filters allow for the creation of blurry yet well-contrasted gratings, the Ricker wavelet (also known as the Mexican hat function) facilitates the design of stimuli with sharper contrasted contours. The rationale behind having two different designs is to assess which properties (blurry vs. sharp) offer the best trade-off between accuracy and visual comfort. We then propose to animate these StAR stimuli using aperiodic short bursts of visual flashes [20]. This method has revealed distinct Visual Evoked Potentials (VEPs) with double the amplitude of classical maximum-length sequences, one of the most employed approaches in c-VEP paradigms (for a review, see [32]). This streamlined approach allowed for rapid calibration, yet maintained high accuracy in classification.

To achieve this goal, we employed a factorial design wherein participants focused on five flickering targets presented in three formats: traditional *Plain* flicker, *Gabor*-based, and *Ricker*-based texture flicker. We conducted subjective assessments to evaluate various dimensions of user experience such as visual comfort, visual tiredness, and peripheral distraction. To demonstrate that our StAR stimuli effectively enhance brain response and facilitate its decoding, we utilized a dry electrode system. Since such systems are known for their lower signal-to-noise ratio compared to wet-based systems [33], reporting high accuracy would validate the robustness of our approach. Consequently, we reported various metrics of BCI performance, including classification accuracy, selection time, and Information Transfer Rate (ITR). Additionally, to provide a comprehensive analysis, we examined evoked responses and inter-trial coherence (ITC) metrics, which can reveal the distinct brain responses elicited by our StAR stimuli. Finally, we implemented an online proof of concept using the Timeflux framework [34] with the stimuli yielding the highest accuracy. This demonstration aims to underscore the potential of deploying BCI beyond the laboratory, featuring a visually comfortable system paired with gel-free EEG technology.

## 2 Stimulus and Burst c-VEP design

This section describes the visual stimuli and types of codes used in both the offline experiment and the online proof of concept.

### 2.1 Visual Stimuli for Augmented Response (StAR)

The primary innovation of our paper lies in our novel approach to utilizing small randomly oriented *grains* as stimuli. This method encompasses the creation of two distinct types of textures: one based on *Gabor* filter and the other on Ricker wavelet. To generate these stimuli, defined by their specified height and width, our algorithm follows a precise sequence of steps. First, it generates a single patch of the specified type (*Gabor* or *Ricker*). Next, it scatters as many copies of this patch as required by the parameters across the texture space, with each copy assigned a random orientation. Finally, these patches are seamlessly blended with the texture background, which we set to a neutral 50% grey. This blending process is crucial as it ensures that any overlapping patches are smoothly integrated rather than simply superimposed, thus eliminating any potential square contour effects. All the stimuli were displayed as 150px×150px discs (see Figure 2-C), with a 50% luminance grey background, and their amplitude depth was also reduced to 70%, which ensures visual comfort while leveraging the transparency of adjacent stimuli in peripheral vision, similarly to our previous study [20].

#### 2.1.1 Gabor stimuli

The *Gabor*-based textures are built upon the foundation of Gabor filters. In our experiment, we designed *Gabor* patches to feature a single white stripe of a Gabor filter, flanked by two black stripes (see Figure1, left). As pixels diverge from the patch’s center, they seamlessly blend with the background, producing a gradient effect. We rely on the following Gabor filter formula:

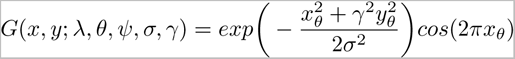

where, *x_θ_* = *x* cos *θ* + *y* sin *θ* and *y_θ_* = −*x* sin *θ* + *y* cos *θ*. Here, *λ* represents the wavelength of the sinusoid, *θ* signifies the orientation of the filter stripes, *ψ* denotes the phase offset of the sinusoid, *σ* represents the standard deviation of the Gaussian envelope, and *γ* characterizes the ellipticity of the support of the Gabor function. To generate our patches (40px×20px), we set the parameters as follows: *λ* = 1, 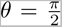, *ψ* = 0, *σ* = 0.8, and *γ* = 1. With these settings, our patch function can be expressed as:

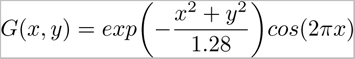

as *x_θ_* and *y_θ_* equal *x* and *y* respectively, with our choice of 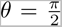. Additionally, we adjusted the background of the selected textures to a medium grey (50% screen luminance). This adjustment serves to mitigate any flickering effects that may occur during the transition to the off-state stimuli color, especially when the texture is fully transparent. Finally, based on test pilot experiments we settled on a total of 75 patches. The left image of Figure 1 shows the Gabor patch detail over a patch-filled texture.

**Fig. 1.**
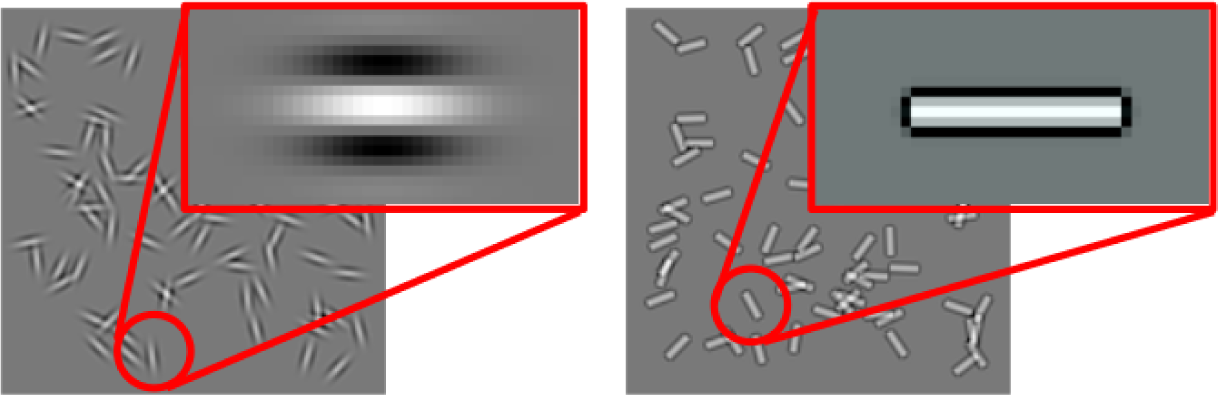
The different types of patches as texture elements. Left: Gabor patch, with its single white stripe surrounded by two black stripes, with the elliptic gradient that blends it into the background. Right: Ricker patch, with its full contour and sharp contrast.

#### 2.1.2 Ricker stimuli

The *Ricker* stimuli are derived from an adaptation of the *Ricker* wavelet function. Unlike the *Gabor* patches, each *Ricker* patch is fully outlined, as opposed to having blurry contours due to the Gaussian gradient applied on the Gabor filter (see Figure1, right). The rationale behind this design is that such patches may offer clearer and more salient visual cues to the receptive fields, thanks to the sharp contrast of the border, potentially enhancing brain response. To create these patches, we utilize the following adaptation of the *Ricker* function:

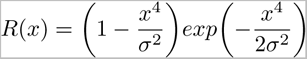

with *σ* = 0.2. These parameters were selected to ensure that the function features a flatter peak at *x* = 0, rather than a sharp peak, thereby ensuring that the white segment of the flicker is sufficiently wide and bordered by a thinner black outline. Using this function, individual patches were created by plotting it in 2D and then stretching it to a predefined ratio. In our experiment, we maintained a width-to-height ratio of 2:1, resulting in the generation of patches with dimensions of 40px×20px. Similarly to the *Gabor* texture, the *Ricker* texture also comprised a total of 75 patches. The right image of Figure 1 shows the Ricker patch detail over a patch-filled texture.

### 2.2 Burst c-VEP

We used five different codes in this study, following a similar approach to our previous work [20]. In that study, we constrained the code generation process to ensure minimal correlation between the codes. One such constraint involved phase-independent minimal correlation, aimed at preventing strong correlations between any two codes at certain relative phases. However, in this experiment, the start and end times of the stimuli presentations are known. Therefore, we can relax this constraint since no code shifting will occur. Unlike in our previous study [20], where we generated numerous codes and selected a subset of the least pairwise correlating ones, we used a different generation approach here. We created a *master code* containing all the burst onsets that would be distributed into the final codes. This ensured that the codes were interleaved, meaning that no two bursts could occur simultaneously, thus enhancing code discriminability. We then derived our codes from this master code using two distribution methods: ordered and pseudo-random.

**Ordered:**

The burst onsets from the *master code* are distributed in the same order to the final codes, i.e. first burst onset to the first code, next one for the second, and so on, cycling through codes in order until all burst onsets have been distributed. This approach allows us to define codes with similar interburst duration, and are visually more periodic.

**Pseudo-random:**

The bursts onset from the *master code* are pseudo-randomly distributed among the codes. The bursts onsets are distributed to the N codes in a random order, then this order is shuffled before distributing the next batch of burst onsets, and so on, until all burst onsets have been distributed. The codes generated with this approach exhibit a higher variance in interburst duration and the codes seem visually less predictable. Finally, we added a constraint that, when changing cycle, the last onset of the previous cycle can not be the same than the first of the next cycle, so as to avoid too short (under 100ms) onset repetition in a single code.

For both the online and offline experiments, the *master code* was a 2.2s long code for a 60Hz display (132 frames), with a minimal 32ms interburst duration (roughly alternating zeroes and ones, with occasionnal 1-frame jitter). This code has been split to accommodate eleven-class experiments, however we only used the first five codes for our experiments. All resulting codes from both methods exhibit 6 burst onsets. The difference then lies in the interburst duration. For the offline experiment, we chose the *ordered* method, resulting in codes with a minimal interburst duration of 350ms, hence a rate of around 3 bursts per second. For the online experiment, we chose the *pseudo-random* method, resulting in codes with intebrurst duration ranging from 8 frames (132ms) to 40 frames (660ms). We aimed to demonstrate the flexibility and validity of both approaches, as code discriminability is maintained through the interleave property and the relatively long interburst duration.

## 3 Offline experiment

### 3.1 Participants

The offline experiment was conducted with twenty four healthy volunteers (4 women, mean age = 29.3 years, SD = 7.5), all students and staff at ISAE-SUPAERO. The research received approval from the ethics committee of the University of Toulouse (CER approval number 2023-749) and was conducted in compliance with the Declaration of Helsinki. Participants provided written informed consent before participating in the experiment. The anonymized data are available at https://zenodo.org/records/11072871.

### 3.2 Experimental Protocol

Participants were seated comfortably and asked to read and sign the informed consent form. The EEG data were collected using the dry 8-electrode Enobio system, with a sampling rate of 500 Hz to capture the surface brain activity. The 8 electrodes were positioned over the occipital and parieto-occipital sites: PO7, O1, O2, O3, PO8, PO3, POz, PO4. EEG data and markers were synchronized during recording using Lab Streaming Layer [35]. Once equipped with the dry 8-electrode EEG system, participants were instructed to direct their attention to five sequentially presented targets, which were cued in a random order for 0.5 seconds each (see Figure 2-B). Subsequently, a 2.2-second stimulation phase ensued, followed by a 0.7-second inter-trial interval. The cue sequence for each trial was pseudo-randomized and varied across blocks. Upon completion of each block, subjects were required to press the space bar to proceed after a brief pause. Participants completed a total of fifteen blocks, with each block consisting of five trials for the three conditions (*Gabor*, *Ricker* or *Plain* stimuli), totaling 75 stimuli per condition (5 targets x 15 blocks), see Figure 2-C. A short video presenting the protocol and the stimuli can be downloaded at the following link https://nextcloud.isae.fr/index.php/s/NQMkSpayWAZfcXb.

**Fig. 2.**
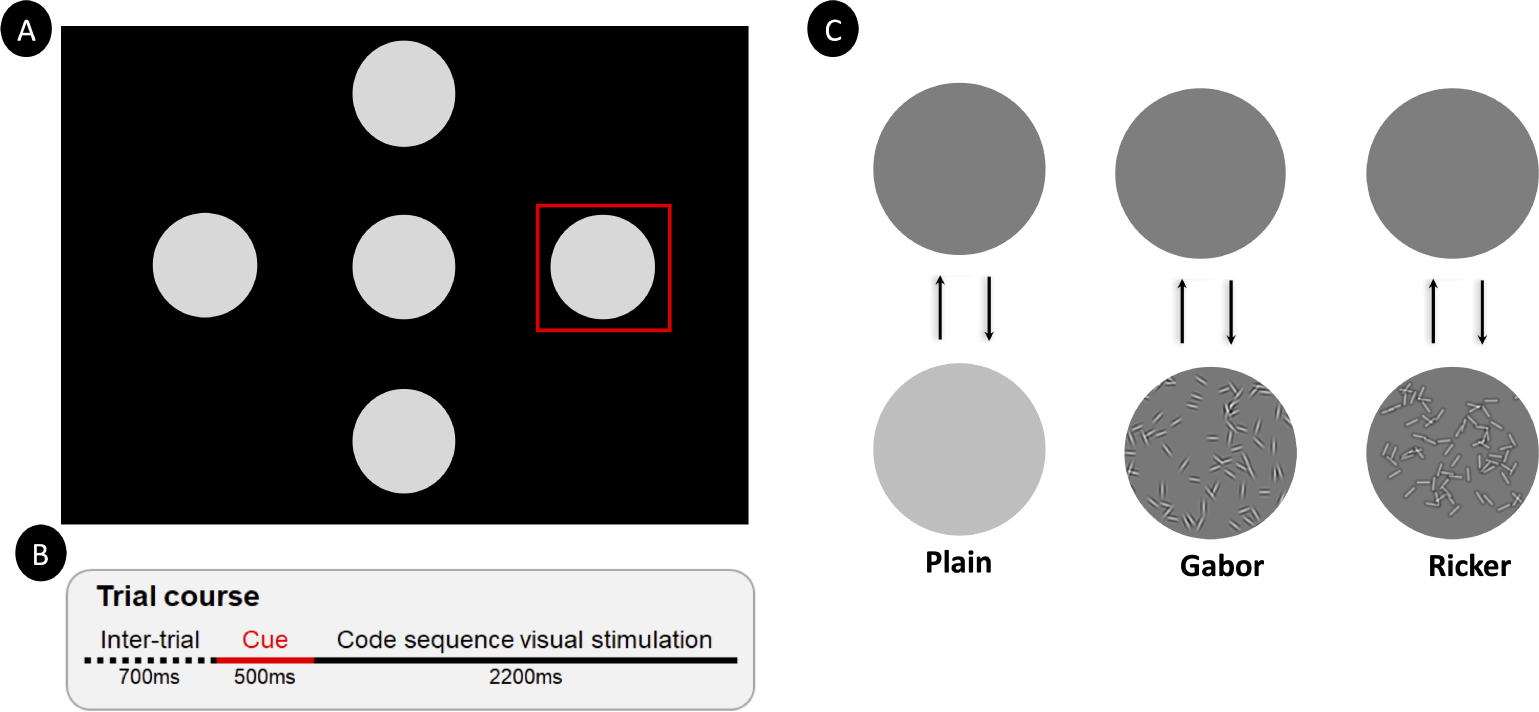
A: The five discs, each of a diameter of 150 pixels, are displayed on an LCD monitor (24.5 inches, 1920 x 1080 resolution, 400*cd/m*^2^, 120 Hz refresh rate, but set to 60 Hz), were presented without borders. A red square served as a cue to direct participants’ attention toward one of the visual stimuli. B: Time course of a trial, an inter-trial interval of 700ms separates the current trial from the previous one. A red-bordered square cue then appears for 500ms around the stimulus to be attended. The cue is followed by the onset of visual stimulation is presented for 2200ms. C: Representation of the possible alternating states for the *Plain*, *Gabor* and *Ricker* stimuli.

The experiment was designed and executed using the Psychopy toolbox in Python ^1^. The stimuli were presented on the following LCD 24.5” monitor: Iiyama Gold Phoenix G-Master GB2590HSU-B1, 1920 × 1080 pixels, 400*cd/m*^2^, and the refresh rate has been set to 60Hz. After each condition, the participants were asked to provide subjective ratings with three 11-points semantic differential scales to rate their subjective experience related to the stimuli presented, including visual comfort (specifically the level of eye strain), mental fatigue (the extent of fatigue induced by the flickers), and peripheral distraction (the visibility and level of distraction caused by peripheral flickers).

### 3.3 VEP Analysis

The effect of experimental manipulations VEP responses was investigated to gain a deeper understanding of the neural mechanisms that could account for variations in classification performance. All the EEG analyses were ran using Matlab (R2021b) [36] and EEGlab (2019.1) [37]. For this purpose, the continuous EEG data was bandpass filtered between 1Hz and 25Hz (FIR, 16501 filter order, cutoff frequencies at 0.05Hz and 25.05Hz). The onset of singular visual stimuli (within the whole sequence) was defined based on the timings at which the code sequences alternated from dark to bright states. The continuous EEG data were then segmented into epochs around the onset of visual stimulation onset (from -0.5s to 1s). The VEP responses were averaged across trials for each electrode and participant. We computed the average VEP response across trials. Subsequently, we determined the mean amplitude of the VEP within a 60ms to 110ms time window and its peak latency from the resulting waveforms at electrode Oz, where the VEP response is most pronounced.

### 3.4 Offline classification of EEG data

The raw continuous EEG data were filtered between 1Hz and 25Hz and re-referenced on POz electrode. The raw data were then epoched from 0 to 250ms relative to flickers onset timestamps. Lastly, a baseline correction was applied to the epochs to eliminate any potential slow drifts. These preprocessed data were then fed into the classificaction algorithm. The *scikit-learn* Python libraries was used to implementating the processing pipeline. The *Pyriemann* library was utilized to transform multivariate time series EEG data into covariance matrices, enabling the classification of the Riemannian geometry of symmetric positive definite matrices. First, the number of trials per class was balanced through the random selection of undersampled ensembles of trials. Then, the dimensionality of the multichannel EEG data was reduced using the unsupervised xDAWN method with a set of four spatial filters [38]. This approach allows the xDAWN algorithm to enhance the distinction between VEP responses from different classes, such as target versus (”1” - burst) non-target VEP (”0”- no burst). Following this, the spatially filtered signals were projected into the estimated signal subspace, generating feature vectors that were then fed into a logistic regression model [39]. To that end, we employed Riemannian geometry methods [40], which have shown superior performance in both single-trial VEP and passive BCI classification tasks, as indicated by a recent retrospective analysis [41]. The BCI pipeline utilized in this study can be formally outlined as follows.

Firstly, a classifier was trained for each subject. A prototype response *P* for each class was obtained by averaging the signals across trials. For each trial *X_i_*, a template trial *T_i_* was created by concatenating *P* with the trial *X_i_*. The template trials *T_i_* were subsequently converted into covariance matrices to capture the spatial characteristics of the signal, as elaborated in [42]. Next, these covariance matrices were projected onto their tangent space, with the geometric mean of all covariance matrices serving as the reference point. Following this projection, each covariance matrix was represented as a vector. Logistic regression, without any regularization, was then applied for classification. Performance was evaluated in terms of accuracy using 5-fold cross-validation, with stratified folds to ensure an equal number of trials for each class in each fold.

Then, the decoding process for predicting the target the subject wants to select is divided into two phases. In the first phase, following the approach of [20, 43], short 250ms windows of EEG data with 2ms steps are input to the classification pipeline. This step decodes the electrophysiological response, resulting in a sequence of ones (indicating the presence of a VEP in the window) and zeroes. Subsequently, this decoded sequence is compared to the stimulation patterns of the targets using Pearson correlation. However, the actual state switch rate of the simulation is limited by the 60 Hz screen refresh rate. To address this, the decoded sequence is downsampled to match the screen refresh rate using majority voting.

In the second phase, once the downsampled decoded sequence reaches a minimal length of 0.7*s*, a Pearson correlation coefficient is calculated between the downsampled binarized decoded sequence and the various target templates. The label of the sequence with the highest correlation is temporarily considered the prediction. The decoded sequence is then extended with the next stimulation decoding using the subsequent EEG data (i.e., the downsampled next 8 windows), and a new temporary prediction is made. If the system consistently outputs the same temporary prediction 60 times, a closing prediction for the trial is triggered, stopping the computation and initiating the classification for the next trial. In situations where no closing prediction could be made after 2.2 *s*, the system does not issue any command.

### 3.5 Statistical Analyses

Statistical analyses were carried out with JASP 0.18 software [44]. A one-way repeated measure ANOVA, with factors of stimulus type (*Gabor*, *Ricker*, *Plain*) were computed for each subjective metrics (visual comfort, mental tiredness and peripheral distraction). Similar one-way repeated measure ANOVA were computed for each objective metrics (classification accuracy, selection time, and information transfert rate) when considering 8 blocks of calibration data (5 flickers × 2.2s × 8 blocks = 88 *s*). Eventually, we ran the same repeated measure ANOVA to assess the effect of types of stimuli on the amplitude of visual evoked potentials (permutation test corrected with false discovery rate for multiple comparison and *p <* 0.01) and ITC. The Greenhouse-Geisser correction was applied when the assumption of sphericity was violated. The Holm test was used for all post-hoc comparisons. Significance level was set at *p <* 0.05 for all analyses.

### 3.6 Results

#### 3.6.1 Subjective results

In the subsequent sections, we present the outcomes of statistical analyses conducted on subjective ratings pertaining to Visual Comfort, Mental Fatigue, and Peripheral Distraction for various stimuli. These findings are depicted on Figure 3.

**Fig. 3.**
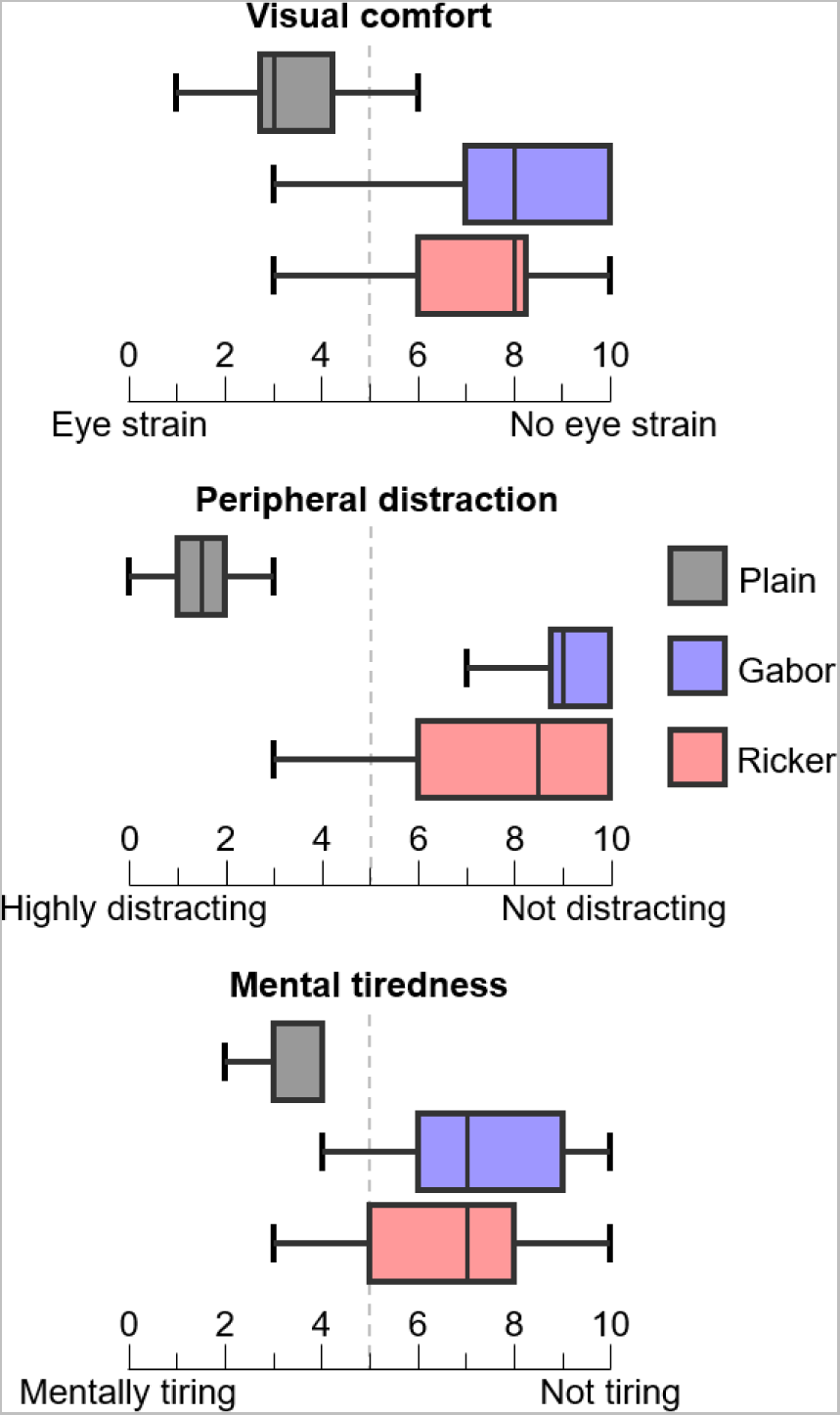
Distribution (N = 24) of subjective assessment scores across three dimensions of user experience for the different types of stimuli (*Plain*, *Gabor*, and *Ricker*). The visual comfort, peripheral distraction, and mental tiredness experienced in relation to the presentation of repeated visual stimuli were measured using a 0-10 semantic differential scales.

##### Visual Comfort

The repeated measure ANOVA disclosed a main effect of stimulus type on visual comfort (F(2,46) = 47.36, *p <* 0.001, 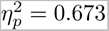). Post-hoc analyses revealed that the *Plain* stimuli (mean = 3.7) were statistically less visually comfortable than the *Gabor* (mean = 8.0, *p <* 0.001, Cohen’s d = -2.081) and *Ricker* (mean = 7.25, *p <* 0.001, Cohen’s d = -1.717). No differences were found between the *Ricker* and the *Gabor* stimuli.

##### Mental Tiredness

The repeated measure ANOVA disclosed a main effect of stimulus type on mental tiredness (F(2,46) = 31.253, *p <* 0.001, 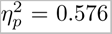). Post-hoc analyses revealed that the *Plain* stimuli (mean = 3.79) led to statistically higher mental tiredness than the *Gabor* (mean = 7.25, *p <* 0.001, Cohen’s d = -1.728) and *Ricker* (mean = 6.79, *p <* 0.001, Cohen’s d = -1.499). No differences were found between the *Ricker* and the *Gabor* stimuli.

##### Peripheral Distraction

The repeated measure ANOVA disclosed a main effect of stimulus type on peripheral distraction (F(2,46) = 76.116, *p <* 0.001, 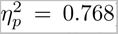). Post-hoc analyses revealed that the *Plain* stimuli (mean = 2.0) were statistically less transparent than the *Gabor* (mean = 8.71, *p <* 0.001, Cohen’s d = -3.077) and *Ricker* (mean = 7.71, *p <* 0.001, Cohen’s d = -2.618). No differences were found between the *Ricker* and the *Gabor* stimuli.

#### 3.6.2 Visual Evoked Potentials analyses

The subsequent sections detail the analyses of evoked responses in both time and frequency domains to aperiodic stimulation across the three different stimulus conditions. Figure 4 summarizes the main findings.

**Fig. 4.**
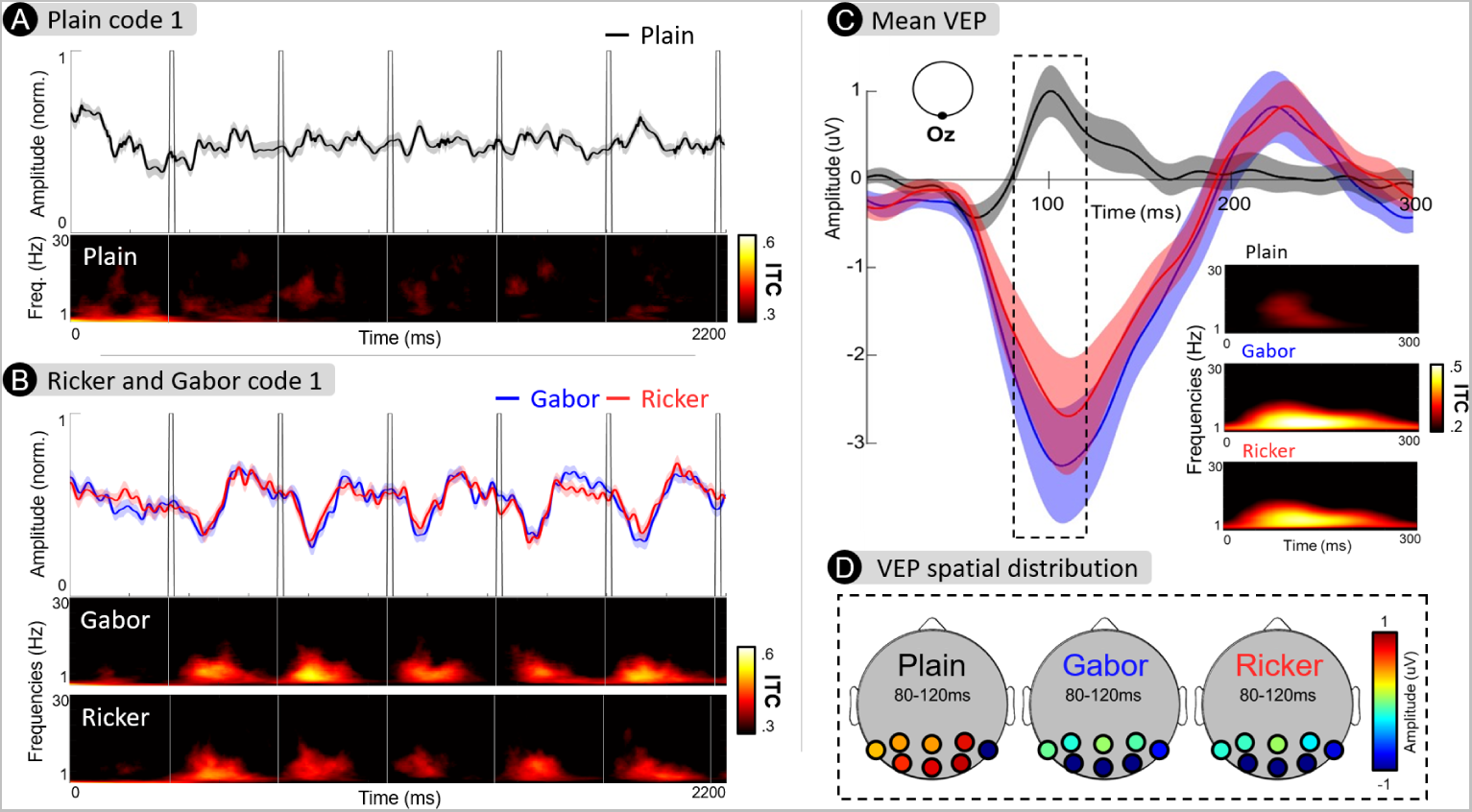
A: Grand average (N=24) of the EEG responses rmeasre at Oz site. The grand average of the Visually Evoked Potentials (VEP) response to the initial burst code-VEP sequence, lasting 2.2 *s*, is shown by the black line. The VEP amplitude has been normalized, and the shaded area represents the standard error of the VEP amplitude across individuals, calculated over time samples. Scalp maps illustrate the instantaneous topographical distribution of VEP responses at different phases of the stimuli input function. The bottom subplot depicts the Inter-Trial Coherence (ITC) of VEPs, computed across frequencies (y-axis) and over time (x-axis), with brighter shades indicating higher ITC. B: Similarly in this subplot, the grand average VEP responses to the initial code-VEP for the *Gabor* and *Ricker* stimuli are presented by the blue and red lines, respectively. C: Grand average (N=24) Visually Evoked Potentials (VEPs) waveforms recorded at electrode Oz for each visual stimuli texture type (*Plain* in black, *Ricker* in red, and *Gabor* in blue). The shaded areas represent standard error around mean. The time-frequency plots in the center present the Inter Trial Coherence for each stimulus type. D: Topographical distribution of VEP responses across the occipital electrodes montage for the 80-120 time window.

**Fig. 5.**
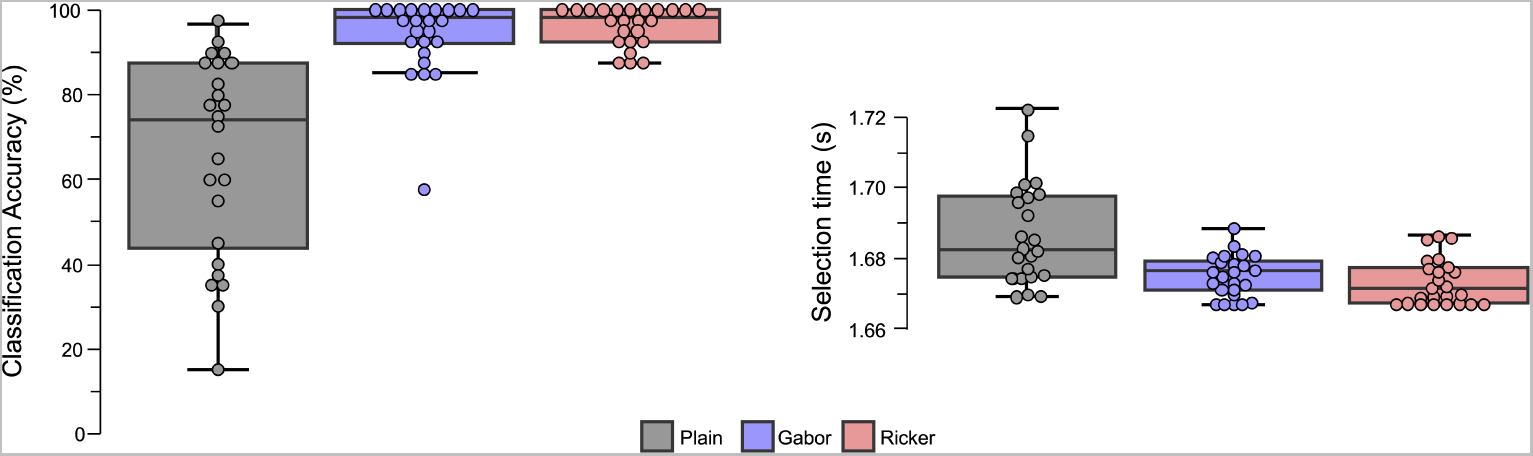
Boxplots presenting the distribution (N = 24) of the offline BCI performance in terms of classification accuracy and selection time for the different types of stimuli (*Plain*, *Gabor*, and *Ricker*). The classification accuracy for each subject is represented by the circles. The median and mean classification accuracy across subjects for each condition is represented by the plain and dotted lines respectively.

##### Amplitude

The repeated measure ANOVA revealed a main effect of stimulus type on VEP amplitude (F(2,46) = 12.66, *p <* 0.001, 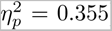). The *Plain* stimuli (mean = 0.93*µV*) exhibited significantly lower VEPs amplitude than the *Gabor* (mean = 2.87*µV*, *p <* 0.001, Cohen’s d = -0.852) and the *Ricker* (mean = 2.81*µV*, *p <* 0.001, Cohen’s d = -0.672) ones. No differences were found between the *Ricker* and the *Gabor* stimuli.

##### Inter-Trial Coherence

The repeated measure ANOVA revealed a main effect of stimulus type on theta ITC (F(2,46) = 8.21, *p <* 0.001, 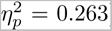). The *Plain* stimuli (mean = 0.18) exhibited significantly lower VEPs amplitude than the *Gabor* (mean = 0.33, *p* = 0.001, Cohen’s d = -0.722) and the *Ricker* (mean = 0.31, *p* = 0.005, Cohen’s d = -0.619) ones. No differences were found between the *Ricker* and the *Gabor* stimuli.

#### 3.6.3 Offline BCI Performances

The following sections provide insights into various performance metrics, including accuracy, decoding time, and ITR.

##### Classification accuracy

The repeated measure ANOVA disclosed a main effect of stimulus type on accuracy (F(2,46) = 45.02, *p <* 0.001, 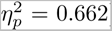). Post-hoc analyses disclosed that the *Plain* stimuli (mean = 65.6%) led to lower accuracy than the *Gabor* (mean = 93.6%, *p <* 0.001, Cohen’s d = -1.869) and the *Ricker* (mean = 96.3%, *p <* 0.001, Cohen’s d = -2.042) stimuli. No differences were found between the *Ricker* and the *Gabor* stimuli.

##### Decoding time

The repeated measure ANOVA disclosed a main effect of stimulus type on decoding time (F(2,46) = 17.290, *p <* 0.001, 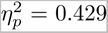). Post-hoc analyses disclosed that the *Plain* stimuli (mean = 1.686*s*) led to slightly slower decoding time than the *Gabor* (mean = 1.675*s*, *p <* 0.001, Cohen’s d = 1.187) and the *Ricker* (mean = 1.673*s*, *p <* 0.001, Cohen’s d = 1.382) stimuli. No differences were found between the *Ricker* and the *Gabor* stimuli.

##### Information Transfer Rate

The repeated measure ANOVA disclosed a main effect of stimulus type on ITR (F(2,46) = 93.435, *p <* 0.001, 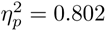). Post-hoc analyses disclosed that the *Plain* stimuli (mean = 31.78) led to lower ITR than the *Gabor* (mean = 69.35, *p <* 0.001, Cohen’s d = -2.132) and the *Ricker* (mean = 73.88, *p <* 0.001, Cohen’s d = -2.389) stimuli. No differences were found between the *Ricker* and the *Gabor* stimuli.

## 4 Proof of concepts : online c-VEP BCI with Ricker stimuli

We implemented an online version of the BCI in which we maintained the identical five-class layout from the offline phase. We selected the *Ricker* stimuli for our experiments due to their superior accuracy and positive user experience ratings.

### 4.1 Participants

The online experiment was conducted with twelve healthy volunteers (2 women, mean age = 29.6 years, SD = 9.6), all students and staff at ISAE-SUPAERO. None of the participants reported any of the exclusion criteria (neurological antecedents, being under psychoactive medication at the time of the study) and had normal or corrected-to-normal vision. Participants gave informed written consent prior to the experiment. The anonymized data are available at https://zenodo.org/records/11072871.

### 4.2 BCI implementation

The online variant was implemented using Timeflux [34], an open-source Python framework dedicated to the real-time acquisition, processing, and classification of time series, with a focus on biosignals for developing fully operational BCI applications through a simple YAML syntax ^2^. Timeflux operates under the principles of Directed Acyclic Graphs (DAG), where data and event streams flow directionally through computing nodes. Multiple graphs can run concurrently, each containing an arbitrary number of nodes connected by edges. The software architecture of the online variant includes six parallel graphs. The *EEG* graph handles data acquisition using the LSL protocol. The *Preprocessing* graph re-references and filters the EEG data. The *Classification* graph constructs and accumulates epochs triggered by the *UI* graph, trains a Machine Learning model, and produces predictions when a defined confidence level is reached. The *UI* graph manages user interaction, providing a conventional monitoring interface for real-time EEG data observation and a specialized BCI interface. The *Record* graph safely stores raw and filtered data and events for further analysis. The *Broker* graph, containing only one node, handles all asynchronous communication (see Figure 6).

**Fig. 6.**
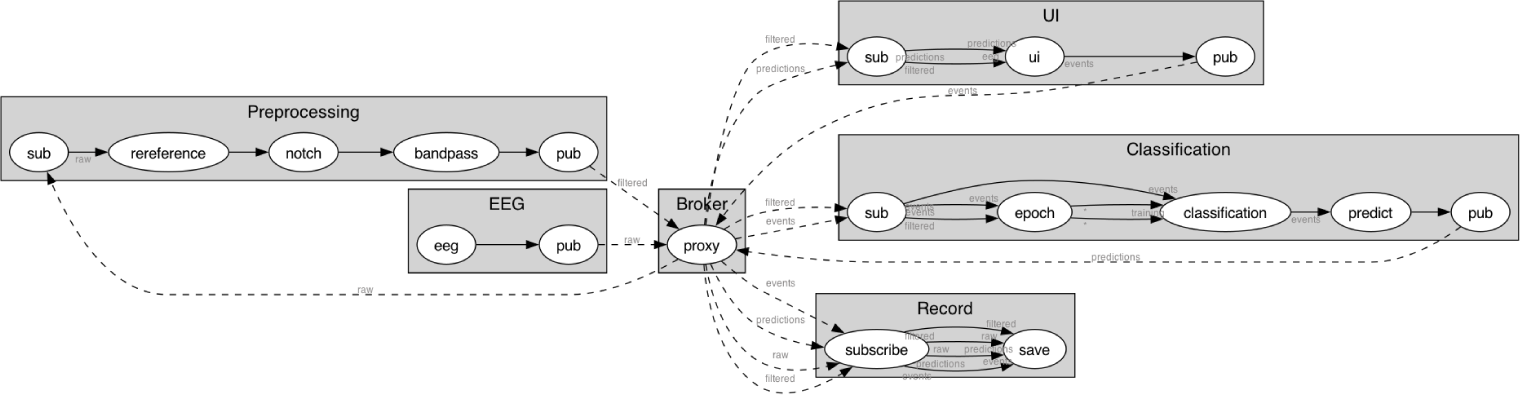
The software architecture of the online implementation, consisting of six graphs running in parallel. Nodes inside each graph are executed in a sequential manner. Data and event streams are represented with dashed lines, and are routed via the special *Broker* graph, according to the asynchronous *publish-subscribe* messaging pattern.

We followed the same preprocessing steps and classification pipeline as in the offline experiment. Epochs were extracted from each frame, synchronized with the screen refresh rate, and sent to the classification engine in near real-time. Predictions were accumulated, and a final prediction was emitted when a configurable threshold was reached. The presentation of stimuli during the different phases (calibration, cued, and digicode tasks) was achieved using Timeflux’s featured JavaScript API in a web browser. For reproducibility purposes, the code and documentation are available ^3^ under a permissive open-source license. The user interface is fully configurable and easily extensible to other task types.

### 4.3 Protocol

During a 66*s* calibration phase (10 random targets among the five, exhibiting 3 repetitions of their 2.2*s* code), participants were exposed to multiple presentations of each c-VEP burst sequence. Subsequently, they engaged in an online cued task, where cues indicated which of the five targets to focus on. Correct predictions resulted in the target turning green, while incorrect ones turned it red, with a total of 20 targets presented. The experiment also included a five-class digital code task, where participants mentally input a sequence of four randomly selected digits (between 1 and 5) displayed on the lower left corner of the screen. Correct predictions by the algorithm turned the target green, allowing participants to proceed to the next digit. Incorrect predictions turned the target red, necessitating a refocus on the target for correct selection. Participants needed to accurately complete five sequences of four digits each, testing the system’s predictive accuracy and the user’s ability to control their focus within this interactive online framework.

### 4.4 Online BCI Performances

The mean classification accuracy for the cued task was 97.5%(*SD* = 3.4) and 94.3% (*SD* = 7.9) for the asynchronous digicode task. The mean decoding time for the cued task was 1.64s (*SD* = 0.12) and was 1.65s (*SD* = 0.09) for the digicode task. The results for each of the 12 participants are presented in Table 1.

**Table 1.**
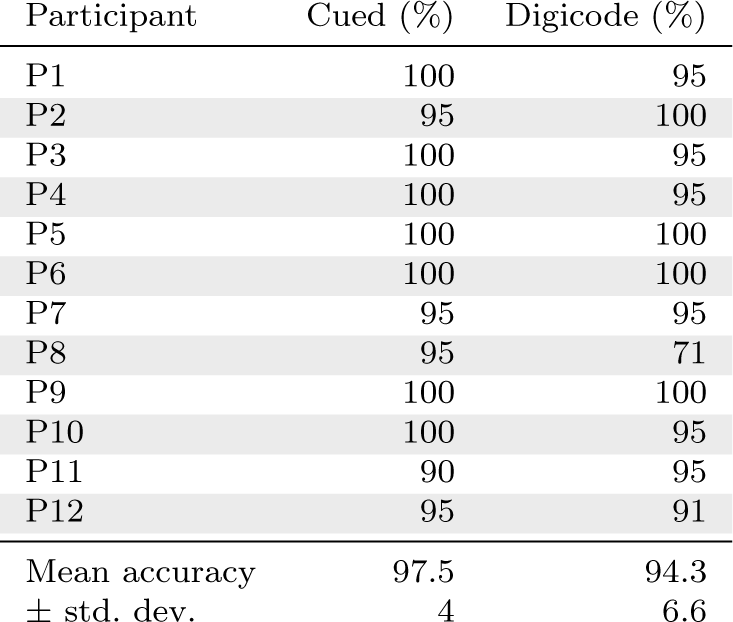
Mean classification accuracy for the cued and asynchronous digicode tasks.

| Participant | Cued (%) | Digicode (%) |
| --- | --- | --- |
| P1 | 100 | 95 |
| P2 | 95 | 100 |
| P3 | 100 | 95 |
| P4 | 100 | 95 |
| P5 | 100 | 100 |
| P6 | 100 | 100 |
| P7 | 95 | 95 |
| P8 | 95 | 71 |
| P9 | 100 | 100 |
| P10 | 100 | 95 |
| P11 | 90 | 95 |
| P12 | 95 | 91 |
| Mean accuracy | 97.5 | 94.3 |
| ± std. dev. | 4 | 6.6 |

## 5 Discussion

The current study focused on the development of innovative RVS stimuli for reactive BCI with the aim of improving visual comfort and peripheral distraction while ensuring effective decoding of neural responses originating from visual cortical areas using dry surface EEG electrodes. Specifically, these stimuli were designed to incorporate small, randomly oriented *Gabor* or *Ricker* patches to optimize their efficacy. To assess the effectiveness of these newly proposed stimuli, we conducted an initial experiment comparing them to the traditionally employed *Plain* flickers. This experiment involved collecting subjective feedback to evaluate the impact of the patches design on user experience, along with offline analyses to determine BCI performance metrics. The insights gained from these evaluations led to the development of an online version of the tasks, demonstrating the practical application and potential benefits of the stimuli. The subsequent sections are dedicated to discussing the principal findings.

### 5.1 End-user assessment

We first propose to examine the subjective findings. The consideration of end-users, particularly in the design of visually comfortable stimuli, has gained significant attention in recent years [15, 19–22, 45]. In our study, we presented textured stimuli that allowed to diminish the luminance since fewer pixels are flickered compared to classical *Plain* flickers. Unsurprisingly, the StAR stimuli, integrating small *Gabor* or *Ricker* patches, were rated as inducing statistically higher visual comfort and lower mental tiredness compared to *Plain* stimuli. More interestingly, the StAR stimuli were not only more imperceptible in peripheral vision than *Plain* stimuli, displaying a statistically significant strong size effect, but the ratings for the *Gabor* and *Ricker* variants were very close to the maximum value (i.e., 10 = no peripheral distraction), reaching mean scores of 8.71 and 7.71, respectively. While one could expect that the *Gabor* stimuli, exhibiting more blurry edges than the *Ricker* stimuli, could be rated as more visually confortable, no statistically significant difference was observed. However, it is worth noting that the variance for the *Ricker* stimuli was much larger than for the *Gabor* ones deviation. These results appear to endorse the effectiveness of employing small gratings to diminish the perception of flickering stimuli in the periphery.

### 5.2 Offline BCI performance

#### 5.2.1 Classification accuracy

While the subjective assessments disclosed that StAR stimuli exhibited crucial end-user features, it was essential to verify their ability to maintain high BCI performance. Typically, one can expect that reducing illumination and stimuli saliency should compromise classification accuracy. However, our classification results not only validated the design intuition behind the StAR stimuli but also surpassed the findings from [46, 47] utilizing c-VEP with dry EEG systems, who reported accuracy rates of 70.1% and 76%. Indeed with only 88*s* of training data, the accuracy rates for the *Ricker* and *Gabor* stimuli reached an 96.3% and 93.6% accuracy, respectively, outperforming the accuracy of *Plain* stimuli at 65.6%. Notably, the standard deviation for the *Plain* stimuli exceeded 20% showing high variability among participants. It is worth noting that in our previous study [20], the same *Plain* stimuli using burst c-VEP reached 95.6% but with a wet research grade EEG system. The significant difference in accuracy with the *Plain* stimuli clearly underscores the contribution of the StAR stimuli in achieving high accuracy with dry EEG systems, known for displaying lower accuracy compared to wet systems. The present results from our BCI study conducted with dry EEG compare favorably with other c-VEP studies that used wet EEG systems, achieving 95.9% accuracy with 384 seconds of calibration data [43], and are consistent with studies like [48], which achieved an average accuracy of 95% in 68*s*, and [49], which achieved 92% accuracy in 84*s*.

#### 5.2.2 Selection time

In line with the accuracy results, both the *Ricker* and *Gabor* stimuli led to slightly faster decoding times compared to the *Plain* stimuli. The decoding time of 1.68 seconds aligns well with previous studies conducted with gel-based electrodes, such as [43] (1.7 seconds), although it is slightly slower than our own study [20] (1.5 seconds). Notably, the selection times achieved through our decoding approach are significantly shorter than those reported with the canonical correlation analysis approach for which the best performances ranged between 3.1*s* and 3.8*s* [15, 16, 50–52]. The brief selection time observed in our study not only contributes to a more user-friendly experience but also enhances the practical applicability of BCI in operational settings.

#### 5.2.3 Electrophysiological evidences

The VEP analyses conducted over electrophysiological signals recorded using only few dry electrodes positioned over the occipital cortex provide a compelling explanation for the superior classification performance observed with patterned stimuli compared to *Plain* ones. Both types of grating stimuli effectively elicited stronger event-related responses, that are inherently more distinguishable, thereby enhancing the machine learning and training process. Consequently, this facilitated the machine learning and training process by providing more reliable and predictable inputs to the classification algorithms. Specifically, the evoked responses (N100-P200) to *Gabor* and *Ricker* stimuli were notably more complex than those to *Plain* stimuli (P100). This heightened complexity can be attributed to the structured nature of *Gabor* or *Ricker* - based gratings, which contain multiple visual features such as spatial frequency and orientation. These patterns involve a broader network of neural populations, eliciting more complex neural responses that lead to the generation of pronounced N100 and P200 responses [53], compared to simpler stimuli, which predominantly result in a P100, as we observed in our previous study [20].

Furthermore, we observed that the early component elicited by patterned stimuli (N100) exhibited an absolute amplitude that was twice as significant as that elicited by *Plain* stimuli (P100). Consistently, the ITC evoked by grating stimuli demonstrated significantly stronger and more distinguishable responses compared to *Plain* stimuli. The *Gabor* and *Ricker* stimuli thus effectively elicited consistent and coherent event-related responses over time in response to our code-VEP sequences (see Figure 4 A and B), thereby providing more reliable and predictable inputs to the classification algorithm. This phenomenon likely stems from the ability of patterned flickers to effectively align with the spatial and temporal frequency preferences of neurons within the visual cortex leading to the generation of more robust VEPs. Additionally, the characteristics of our patterned flickers, which were less perceptible due to their small size and low luminance, might have necessitated heightened attentional engagement [54].

### 5.3 Online BCI

We then performed an online implementation of the BCI to demonstrate the effectiveness of our approach. To validate the online framework, participants first engaged in a cued task that mirrored the setup of the initial experiment, allowing for a direct comparison. The accuracy of the cued online task modestly surpassed that of the offline task, despite the decoding times being similarly brief. This variance is likely due to the reduced sample size of our study, which was half that of the offline experiment, coupled with a low number of targets presented to the participants. Our participants were then engaged in a second, self-paced digicode task. Despite a slight reduction in accuracy, the results remained high. This dip in performance can be chiefly attributed to two factors: firstly, the asynchronous nature of the digicode task eliminated any cues indicating which target to focus on, preventing participants from preparing and fixating on a target. Secondly, the task introduced a heightened level of complexity, necessitating the execution of back-and-forth eye movements between the instructed sequence and the target and involved working memory. Previous studies indicated that task complexity can affect BCI performance [55, 56]. The online results affirm that all participants successfully operated the BCI, as evidenced by minimal subject standard deviation and the absence of BCI literacy issues.

Going beyond the implementation of visually comfortable stimuli, our c-VEP BCI design, using brief aperiodic visual flashes, address several end user experience issues in BCI such as calibration time, flexibility and sense of control. While some authors implemented calibration free BCI [52], generally, this result with significantly longer decoding time (e.g., 5s) which in turn may frustrate the users. As usual, a good trade-off have to be found between accuracy, BCI reactivity and calibration time. While straightforward, our accumulative decision based process also facilitates the integration of early stopping mechanisms by enabling decisions to be made without waiting for the entire cycle of the code sequence, as required in template-based approaches [32]. Furthermore, our BCI enables an asynchronous approach, freeing users from the constraints of synchronizing with predetermined stimulation pacing. This feature liberates users from imposed command selection paces, thus preserving their interaction freedom with the interface. Finally, our demonstration of compatibility with gel-free EEG technology underscores the versatility and practicality of our BCI system.

## 6 Conclusion

The findings of this study highlight the transformative potential of StAR stimuli when coupled with the *burst* code VEP paradigm, driving the evolution of BCI technology beyond the confines of the laboratory. Our implementation achieved high accuracy levels with a dry EEG system, requiring only minimal calibration data. This study underscores the paramount importance of integrating visual neuroscience and addressing end-user-related concerns in stimulus design. These stimuli not only enhance visual comfort but also optimize cerebral response over the occipital cortex, facilitating decoding with a dry EEG system. This outcome holds significant implications not only for the BCI community but also for broader cognitive neuroscience research. Our StAR stimuli, characterized by comfort and subtle perceptibility in the periphery, possess the potential for application in various reactive BCI paradigms such as P300 speller, SSVEP, and visual oddball-based BCI.

## 7 Declarations

- This research was funded by ANITI (Artificial and Natural Institute Toulouse Institute) and the AXA research fund.
- The authors declare no conflict of interest/competing interests
- This study was approved by the ethics committee of the University of Toulouse (CER approval number 2023-749) and was carried in accordance with the declaration of Helsinki.
- The anonymized data are available at https://zenodo.org/records/11072871.
- The code and documentation are available at https://github.com/timeflux/burst/
- Author contribution: F.D.: conceptualization, data collection, data analysis, writing; K.C.C.: conceptualization, implementation, data analysis, writing; S.L., data analysis, writing, P.C.: Timeflux implementation, writing

## Footnotes

1 https://www.psychopy.org/

2 https://doc.timeflux.io

3 https://github.com/timeflux/burst/

